# Ubiquitination of the GluA1 subunit of AMPA receptors is required for synaptic plasticity, memory and cognitive flexibility

**DOI:** 10.1101/2022.07.27.501670

**Authors:** Sumasri Guntupalli, Pojeong Park, Dae Hee Han, Mitchell Ringuet, Daniel G. Blackmore, Dhanisha J. Jhaveri, Frank Koentgen, Jocelyn Widagdo, Bong-Kiun Kaang, Victor Anggono

**Affiliations:** Clem Jones Centre for Ageing Dementia Research, The University of Queensland, Brisbane, Queensland, 4072, Australia; Queensland Brain Institute, The University of Queensland, Brisbane, Queensland, 4072, Australia; School of Biological Sciences, Seoul National University, Seoul, 08826, Korea; Mater Research Institute, The University of Queensland, Brisbane, Queensland, 4072, Australia; Ozgene Pty Ltd., Bentley DC, Western Australia, 6983, Australia

**Keywords:** AMPA receptors, ubiquitination, synaptic plasticity, memory, cognition, knock-in mice

## Abstract

Activity-dependent changes in the number of AMPA-type glutamate receptors (AMPARs) at the synapse underpin the expression of long-term potentiation (LTP) and long-term depression (LTD), cellular correlates of learning and memory. Post-translational ubiquitination has emerged as a key regulator of the trafficking and surface expression of AMPARs, with ubiquitination of the GluA1 subunit at Lys-868 controlling the post-endocytic sorting of the receptors into the late endosome for degradation, and thereby regulating their stability at synapses. However, the physiological significance of GluA1 ubiquitination remains unknown. In this study, we generated mice with a knock-in mutation in the major GluA1 ubiquitination site (K868R) to investigate the role of GluA1 ubiquitination in synaptic plasticity, learning and memory. Our results reveal that these mice have normal basal synaptic transmission but exhibit enhanced LTP and deficits in LTD. They also display deficits in short-term spatial memory and cognitive flexibility. These findings underscore the critical roles of GluA1 ubiquitination for bidirectional synaptic plasticity and cognition.

## INTRODUCTION

Long-lasting changes in synaptic transmission, termed synaptic plasticity, have long been considered a cellular model of learning and memory (Huganir and Nicoll, 2013). Long-term potentiation (LTP) and long-term depression (LTD) are two forms of synaptic plasticity that have been intensively studied in this context. Key molecular events underlying the expression of LTP and LTD involve an increase or decrease in the number of AMPA receptors (AMPARs) at the postsynaptic membrane, respectively (Anggono and Huganir, 2012). GluA1 is one of the predominant subunits expressed in the hippocampus and cortex (Lu et al., 2009). GluA1 knockout mice display profound deficits in hippocampal LTP, and working and spatial memory (Sanderson et al., 2007; Sanderson et al., 2010; Zamanillo et al., 1999), underscoring the importance of GluA1-containing AMPARs in synaptic plasticity and cognition.

The C-terminal domain of GluA1 contains multiple sites for protein-protein interaction and post-translational modifications that are critical in regulating the membrane trafficking of AMPARs (Anggono and Huganir, 2012; Diering and Huganir, 2018; Lu and Roche, 2012). Reversible post-translational modifications such as phosphorylation, palmitoylation and ubiquitination in the GluA1 subunit provide fine-tuning mechanisms that control synaptic expression of the receptors (Diering and Huganir, 2018; Hayashi, 2021; Lussier et al., 2015; Widagdo et al., 2017). The best characterised among these is the phosphorylation of GluA1 at the Ser-831 and Ser-845 residues, which are known to control AMPAR trafficking and gating in neurons (Hu et al., 2007; Kristensen et al., 2011; Man et al., 2007). These phosphorylation sites are tightly regulated during LTP and LTD (Lee et al., 2000). Pseudophosphorylation of GluA1 at Ser-831 and Ser-845 is sufficient to lower the threshold for LTP induction in the GluA1 S831/S845D phosphomimetic knock-in mouse (Makino et al., 2011). Furthermore, GluA1 S831/845A phospho-deficient knock-in mice show deficits in LTP and LTD, as well as memory defects in spatial learning tasks (Lee et al., 2003). This evidence highlights the importance of GluA1 phosphorylation events at these residues for synaptic plasticity, learning and memory.

Post-translational ubiquitination has emerged as an important mechanism that regulates the surface expression of AMPARs (Goo et al., 2015; Mabb, 2021; Widagdo et al., 2015). There is also considerable functional cross-talk between the ubiquitination and phosphorylation of different AMPAR subunits (Guntupalli et al., 2017; Widagdo et al., 2020). Activity-dependent ubiquitination of GluA1 occurs at the C-terminal Lys-868 residue and is primarily mediated by E3 ligases, Nedd4-1 and Nedd4-2 (Lin et al., 2011; Schwarz et al., 2010; Zhu et al., 2017). GluA1 ubiquitination controls the post-endocytic sorting of AMPARs toward late endosomes for degradation (Lin et al., 2011; Schwarz et al., 2010; Widagdo et al., 2015; Widagdo et al., 2017). In pathological conditions, excessive ubiquitination of GluA1 mediates amyloid-β (Aβ)-induced downregulation of surface AMPAR expression, leading to synaptic depression (Guntupalli et al., 2017; Rodrigues et al., 2016; Zhang et al., 2018). However, the physiological functions of GluA1 ubiquitination in synaptic plasticity, learning and memory remain unknown. Here, we report the generation and characterisation of a knock-in mouse harbouring the K868R mutation in the *Gria1* gene that encodes the GluA1 subunit of AMPARs.

## RESULTS

### GluA1 K868R ubiquitin-deficient mice have normal gross brain morphology and AMPAR distribution at synapses

To directly test if GluA1 ubiquitination is necessary for synaptic plasticity, learning and memory, we generated a mutant mouse line that contains an arginine substitution at the major GluA1 ubiquitination site, Lys-868, by targeting the mouse *Gria1* gene using the homologous recombination technique (Figure 1A). Chimera mice carrying the mutant allele were bred to CMV-Cre mice to generate the conventional GluA1 K868R knock-in mice. The success of these procedures was confirmed by PCR analysis and direct DNA sequencing (Figures 1B and 1C). We further confirmed the effect of the mutation on ligand-induced GluA1 ubiquitination in primary cortical neurons derived from wild-type or GluA1 K868R knock-in mice. As expected, a 10 min treatment of AMPA induced an increase in the levels of GluA1 ubiquitination in the wild-type neurons, but not in those derived from GluA1 knock-in mice (Figures 1D and 1E). However, this mutation did not affect AMPA-induced ubiquitination of the GluA2 subunit of AMPARs (Figure S1). The GluA1 ubiquitination mutant mice showed no overt behavioural phenotype and bred normally. At 3 months of age, these knock-in mice had normal body weight (Figure 1F) and did not display any gross brain abnormalities compared to their wild-type littermates. Nissl-stained brain sections from homozygous mice also revealed no significant differences in cytoarchitecture (Figure 1G). Taken together, these results suggest that the lack of GluA1 ubiquitination has no effect on gross brain anatomy.

**Figure 1.**
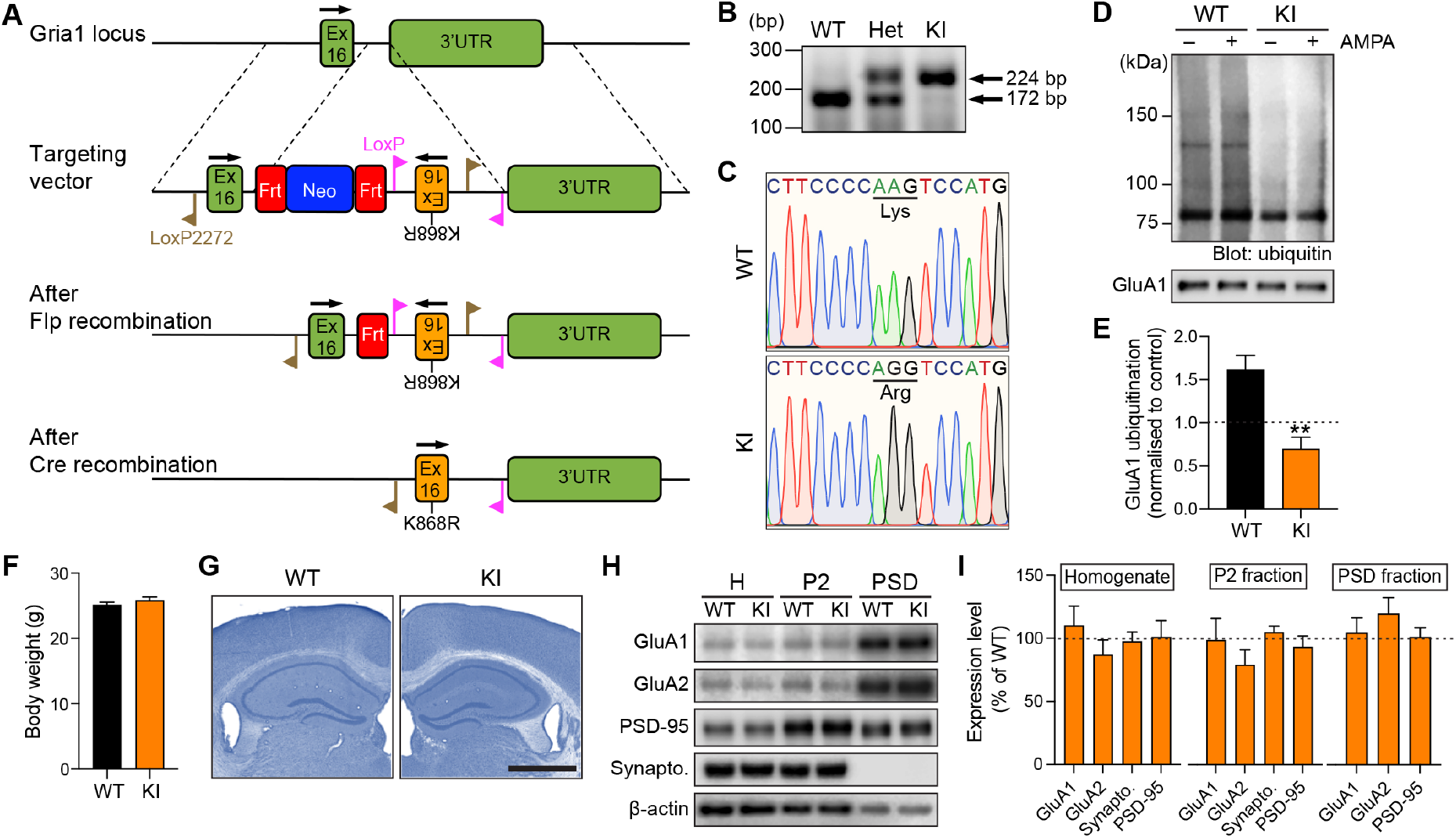
Normal gross anatomy and distribution of GluA1 and GluA2 subunits in the forebrain of GluA1 K868R knock-in mice. (A) Schematic representation of the genomic DNA structure of the *Gria1* locus that encodes the C-terminal tail of the GluA1 protein, the targeting vector, and the resulting genomic structures following Flp- and Cre-mediated recombination events. Ex, exon; UTR, untranslated region; Frt, flippase target site; Neo, neomycin gene. (B) PCR analysis of genomic DNA samples from wild-type (WT), heterozygous (Het) and homozygous (KI) mutant mice. (C) Verification of the GluA1 K868R mutation in homozygous KI mice, and its absence in WT mice by DNA sequencing. (D) Primary cortical neurons derived from WT or GluA1 K868R KI mice were incubated in the presence or absence of 100 μM AMPA, lysed and immunoprecipitated with anti-GluA1 antibodies. Eluted proteins were subjected to western blot analysis and probed with anti-ubiquitin and anti-GluA1 antibodies. (E) Quantification of the levels of AMPA-induced GluA1 ubiquitination normalised to the controls (dashed line). Data are presented as mean ± SEM (n = 5 per group). \*\**p* = 0.0079 using a Mann-Whitney test. (F) No differences were observed in the body weights of GluA1 WT and KI mice. Data are presented as mean ± SEM (n = 15 per group). *p* = 0.49 using a Mann-Whitney test. (G) Nissl-stained hippocampal sections of GluA1 WT and KI mice show normal cytoarchitecture. Scale bar, 1 mm. (H) Whole forebrain lysates (H, total homogenate), crude synaptosomes (P2) and PSD fractions from 3-month-old GluA1 KI and WT mice were immunoblotted with specific antibodies against GluA1, GluA2, synaptophysin, PSD-95 and β-actin. (I) Quantification of the levels of GluA1, GluA2, synaptophysin and PSD-95 in each fraction after normalisation with β-actin. The average protein expression levels in the GluA1 KI mice were normalised to those of WT mice (dashed line). All data are represented as mean ± SEM (n = 7 - 13 per group).

To determine whether the lack of a major GluA1 ubiquitination site changes the distribution of AMPARs at postsynapses, we performed biochemical subcellular fractionation on forebrain tissues of GluA1 knock-in and wild-type mice. Enrichment of the postsynaptic marker PSD-95 and the absence of the presynaptic marker synaptophysin in the postsynaptic density (PSD) fraction demonstrated the purity of this fraction in our preparation. We observed similar levels of GluA1, GluA2, PSD-95 and synaptophysin proteins in the total homogenate, crude synaptosomal (P2) and PSD fractions (Figures 1H and 1I). These data demonstrate that the ubiquitination of GluA1 is not critical for maintaining the steady-state level of synaptic AMPARs under basal conditions *in vivo*, consistent with our previous findings in primary neuronal cultures (Widagdo et al., 2015).

### LTP is enhanced and LTD is reduced in GluA1 K868R knock-in mice

Next, we examined the role of GluA1 ubiquitination on Lys-868 on the expression of LTP and LTD in adult male mice. We first used a single episode of theta-burst stimulation (TBS, 5 bursts at 5 Hz, with each burst composed of five pulses at 100 Hz) to induce LTP, revealing a significant enhancement of input-specific LTP in hippocampal slices from GluA1 knock-in mice compared to their wild-type littermates (Figure S2). Moreover, LTP induced by three episodes of TBS with a 10 min inter-episode interval, which is dependent on the activity of protein kinase A (PKA) and new protein synthesis (Park et al., 2014), was also greatly enhanced in the GluA1 knock-in slices (Figure 2A). In contrast, both NMDA receptor (NMDAR)- and metabotropic glutamate receptor (mGluR)-dependent LTD induced by low-frequency stimulation and bath application of DHPG, respectively, were impaired in GluA1 knock-in slices (Figures 2B and 2C).

**Figure 2.**
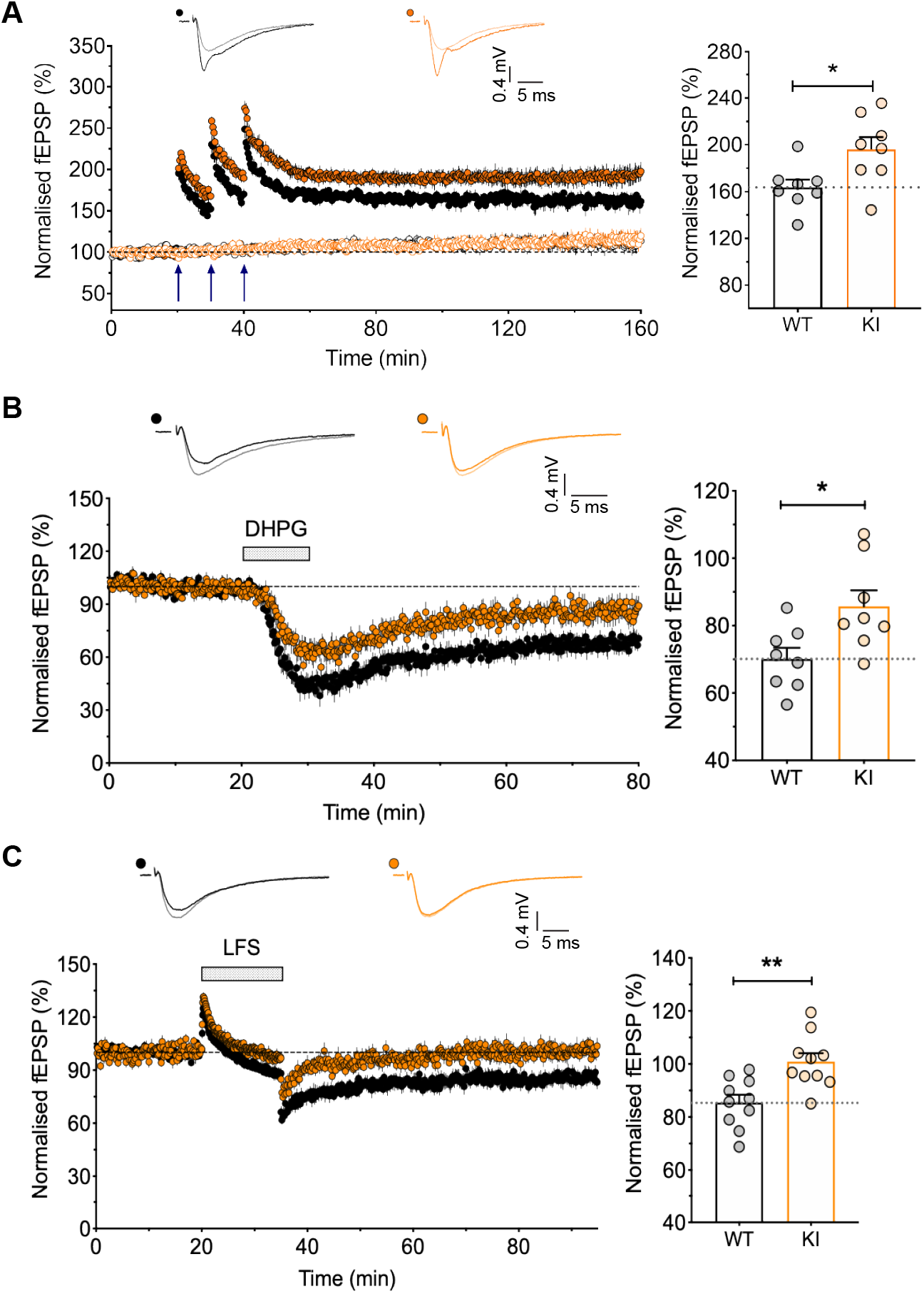
Enhanced LTP and reduced LTD in GluA1 K868R knock-in mice. (A) Input-specific LTP induced by three episodes of theta-burst stimulation with a 10 min inter-episode interval and monitored for 2 h following LTP induction. The right panel shows a significantly larger LTP in the GluA1 KI (n = 8/8) compared to WT littermates (n = 8/7). Data represent mean ± SEM. \**p* = 0.0188 using an unpaired *t*-test. (B) LTD was induced by the bath application of DHPG (100 µM) + D-AP5 (50 µM). The right panel shows the quantification of the levels of LTD. Data represent mean ± SEM (WT, n = 8/8 and KI, n = 8/8). \**p* = 0.017 using an unpaired *t*-test. (C) LTD was induced by low-frequency stimulation (LFS, 1 Hz for 15 min). Data represent mean ± SEM (WT, n = 10/10 and KI, n = 10/10). \*\**p* = 0.002 using an unpaired *t*-test.

To confirm that the enhanced LTP and deficits in LTD in the adult GluA1 ubiquitination-deficient mice were not due to non-specific effects on synaptic transmission, we investigated the basal synaptic properties of hippocampal slices prepared from these mice. Extracellular field potential recordings with varying stimulus intensity generated identical input-output curves from homozygous knock-in mice and their wild-type littermates (Figure 3A). We also found no change in paired-pulse facilitation across different inter-stimulus intervals (Figures 3B). These data suggest that GluA1 ubiquitination does not affect the number of Schaffer collateral-CA1 synaptic connections or short-term presynaptic plasticity. We then performed whole-cell recordings and measured the ratio of NMDAR to AMPAR-mediated synaptic currents and found no statistical differences between the GluA1 knock-in and wild-type neurons (Figure 3C). In addition, the rectification index of AMPARs was also unchanged in GluA1 knock-in neurons (Figure 3D), suggesting that GluA1 ubiquitination does not affect the recruitment of Ca^2+^-permeable GluA1 homomers to the synapse under basal conditions. Furthermore, we observed no significant differences in either the average amplitude or frequency of miniature excitatory postsynaptic currents (mEPSCs) between GluA1 knock-in and wild-type neurons (Figures 3E and 3F). Collectively, these data suggest that alterations in bidirectional synaptic plasticity are not due to abnormalities in basal synaptic transmission or NMDAR-mediated responses, but are likely to be caused by the absence of ubiquitination-dependent regulation of GluA1 function during LTP and LTD.

**Figure 3.**
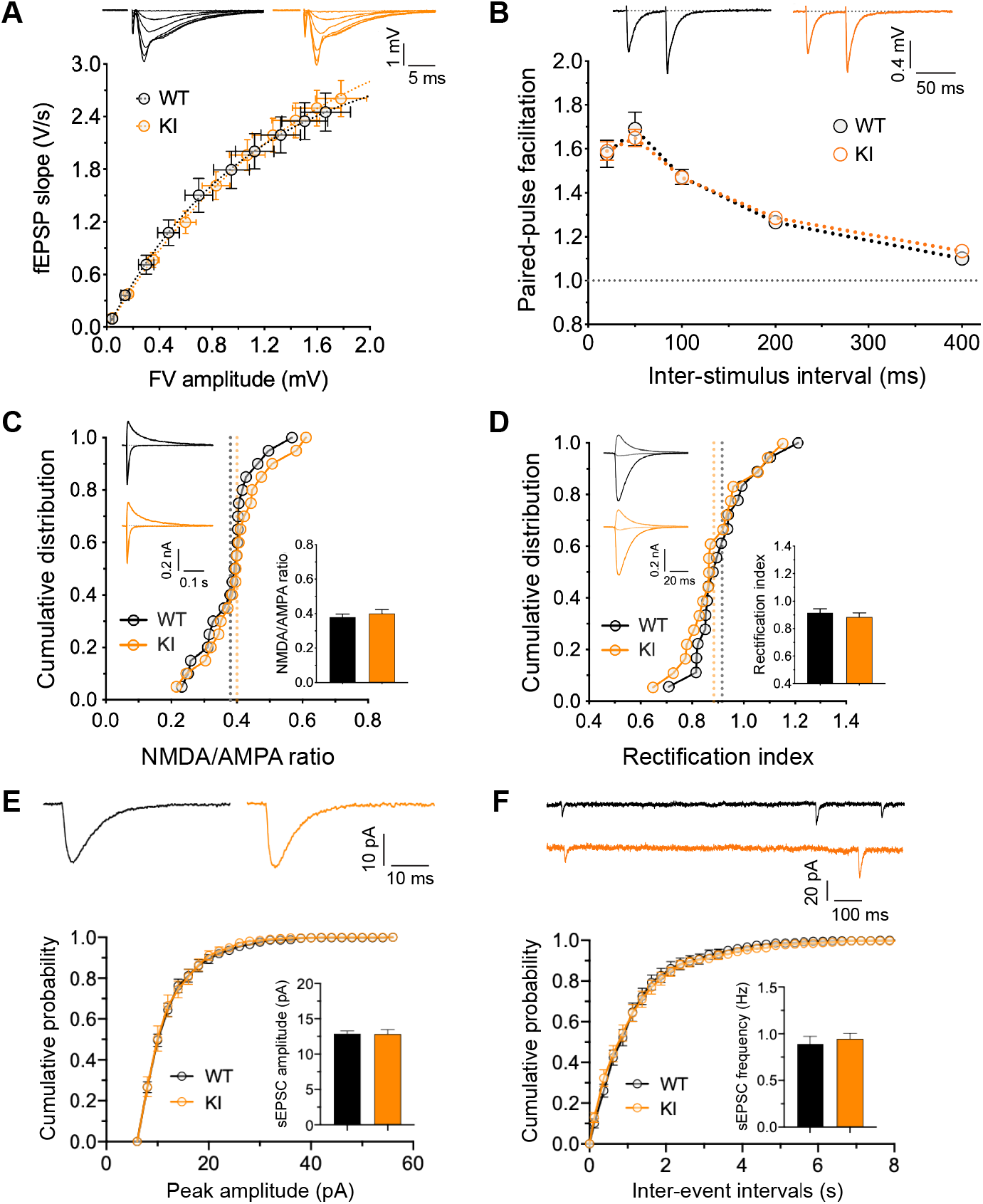
Intact basal synaptic properties in GluA1 K868R mutant mice. (A) Input-output relationships of evoked field EPSP (fEPSP) responses at CA1 synapses for WT (black circles; n = 10 from 10 mice) and GluA1 K868R KI (orange circles; n = 10/10) littermates (male, 2-3 months old). The insets indicate representative traces (averages of 4 successive sweeps). The stimulus intensity is represented by fibre volley (FV) amplitude. (B) Paired-pulse facilitation estimated at inter-stimulus intervals of 20, 50, 100, 200 and 400 ms (WT, n = 10/10 and KI, n = 10/10). (C) NMDA/AMPA ratio measurements in the presence of picrotoxin (50 µM) + (+)-bicuculline (10 µM) for GABA_A_ receptor inhibition by varying holding potentials from -70 mV to +40 mV. Data represent mean ± SEM (WT, n = 20/13 and KI, n = 20/13). *p* = 0.46 using an unpaired *t*-test. (D) AMPAR rectification measured in the presence of D-AP5 (50 µM) + L-689,560 (5 µM) for NMDAR inhibition in addition to the GABA_A_ receptor blockers. AMPAR currents were measured at -70, 0 and +40 mV. Data represent mean ± SEM (WT, n = 18/13 and KI, n = 18/13). *p* = 0.44 using an unpaired *t*-test. (E and F) Amplitude (E) and frequency (F) of miniature EPSC (mEPSC) recordings in the presence of tetrodotoxin (0.5 µM). Data represent mean ± SEM (WT, n = 17/4 and KI, n = 13/5). *p* = 0.945 (E) and *p* = 0.613 (F) using an unpaired *t*-test.

### GluA1 K868R knock-in mice exhibit cognitive deficits

Given the profound alterations in LTP and LTD in the hippocampal CA1 region, we next examined hippocampal-dependent cognitive performance in the GluA1 knock-in mice. First, we used the Y-maze novelty preference test to measure short-term spatial memory, in which mice are allowed to explore the previously unvisited novel (blocked) and previously visited arms (Kraeuter et al., 2019) (Figure 4A). When tested 90 min after the training sessions, both wild-type and GluA1 knock-in mice entered the starting and novel arms with the same frequencies (Figure 4B). However, the wild-type, but not the homozygous mutant mice spent significantly more time in the novel arm (Figure 4C), suggesting that the GluA1 ubiquitin-deficient mice have impaired short-term memory. These mice were also subjected to the elevated plus maze and open field tests to assess their general anxiety-related behaviour and locomotion. No obvious differences in anxiety levels or locomotor activity were observed in GluA1 knock-in mice compared to their wild-type littermates (Figures S3A-S3G).

**Figure 4.**
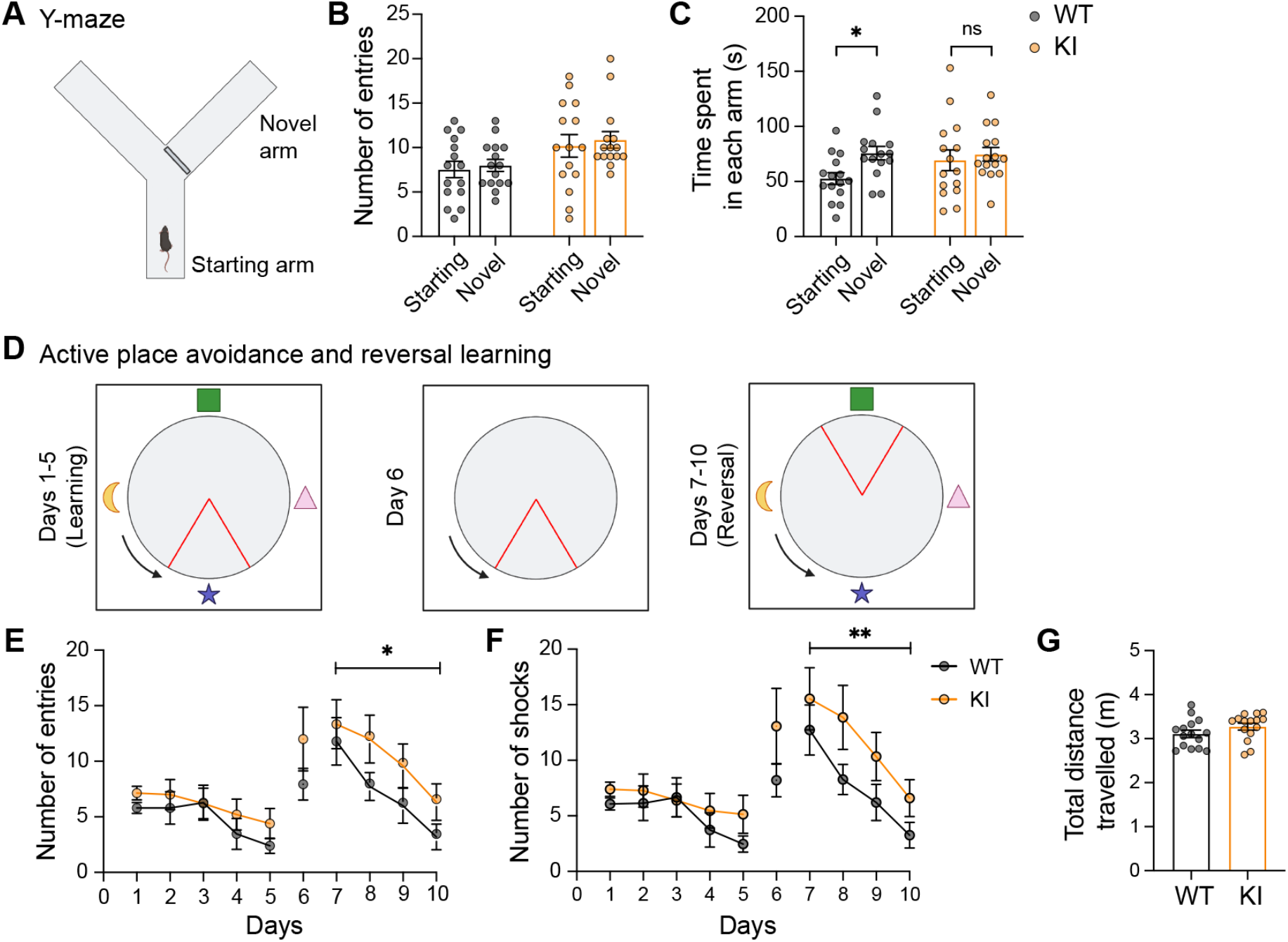
GluA1 K868R mutant mice have cognitive deficits. (A – C) Measurement of short-term spatial working memory using the Y-maze with an initially blocked arm (A). Both WT and GluA1 KI littermates exhibited the same number of entries into the starting and the novel arms when opened (B). However, GluA1 WT, but not the KI mice, spent significantly more time in the novel arm (C). \**p =* 0.044 using a two-way ANOVA with Sidak’s multiple comparison test. (D) Schematic representation of the experimental design for the active place avoidance and reversal learning tasks. The arena contained visual cues on each side of the wall. Blue arrows and red triangles indicate the arena rotation and shock zone location, respectively. (E and F) GluA1 KI mice displayed normal learning during the first 5 days but had significantly more entries into the shock zone (E) and shocks received (F) during reversal learning. F(1, 112) = 6.033, \**p* = 0.016 (E) and F(1, 112) = 7.278, \*\**p* = 0.0081 (F) using a two-way ANOVA test. (G) There was no difference in total distance travelled during the task (Mann-Whitney test, *P* = 0.16). All data are represented as mean ± SEM (*n* = 15 mice per group).

To further examine the cognitive performance of GluA1 knock-in mice, we conducted an active place avoidance test, a well-established hippocampal-dependent spatial memory task (Willis et al., 2017). This experiment was designed to include a reversal-learning task to test the cognitive flexibility of the mice (Figure 4D). Both wild-type and GluA1 knock-in mice exhibited robust learning over 5 days as evidenced by a significant decrease in the number of entries into the shock zone (Figure 4E). However, when the shock zone was moved to the opposite quadrant during reversal learning (days 7-10), GluA1 knock-in mice learnt to avoid the shock zone but committed significantly more errors compared to wild-type mice (Figure 4F). This suggests a specific deficit in cognitive flexibility in the GluA1 mutant mice. No significant difference in total distance travelled during the test period was noted between the genotypes, indicating that locomotion was not affected (Figure 4G).

## DISCUSSION

Upon ligand binding, the Lys-868 residue in the C-terminal tail of GluA1 is covalently modified by K63-linked polyubiquitination, which in turn regulates the post-endocytic sorting of AMPARs towards late endosomes for degradation (Schwarz et al., 2010; Widagdo et al., 2015; Widagdo et al., 2017). However, the physiological importance of GluA1 ubiquitination in neurotransmission, synaptic plasticity, learning and memory remains unknown. In this study, we generated and characterised the GluA1 K868R ubiquitin-deficient knock-in mice. Our analysis revealed that these mice have normal gross brain cytoarchitecture, intact basal synaptic transmission and breed normally. However, the GluA1 mutant mice exhibited enhanced LTP and impaired LTD. Although these mice display normal hippocampal-dependent spatial learning, they have impairments in short-term memory and cognitive flexibility. Our findings therefore demonstrate the critical roles of ubiquitin-mediated post-endocytic sorting and degradation of AMPARs in synaptic plasticity and cognition.

The role of GluA1 ubiquitination in regulating the surface expression and turnover of AMPARs under basal conditions remains controversial (Widagdo et al., 2017). Our current findings are largely in line with our previous data demonstrating that mutation of the GluA1 ubiquitination site does not alter the basal turnover and surface expression of AMPARs in primary neuronal cultures (Widagdo et al., 2015), which are in contrast to those reported in other studies (Lin et al., 2011; Schwarz et al., 2010). Firstly, we did not observe any changes in the protein expression of either GluA1 or GluA2 in the crude synaptosomal and PSD fractions in the forebrain of GluA1 knock-in mice. Secondly, we found that basal synaptic transmission in the CA3-CA1 synapses in the hippocampus of these mice remains intact, with no significant differences in spontaneous or evoked AMPAR-mediated synaptic currents. We also observed no significant changes in the rectification index of AMPARs, indicating that GluA1 ubiquitination does not regulate the subunit composition of AMPARs at the synapse under basal conditions. This makes sense given that ubiquitination of GluA1 occurs at extremely low levels under basal conditions and is only induced upon ligand binding to the AMPARs.

AMPAR recycling is essential for the maintenance of both LTP and LTD (Brown et al., 2007; Fernandez-Monreal et al., 2012; Park et al., 2004; Parkinson and Hanley, 2018; Petrini et al., 2009). Fundamentally, endosomal recycling of AMPAR-containing vesicles provides a constant supply of AMPARs to the plasma membrane in order to maintain LTP, while the sorting of AMPARs into late endosomes for lysosomal degradation leads to a reduction in synaptic strength that is associated with LTD. The fact that GluA1 knock-in mice have enhanced LTP and deficits in LTD is entirely consistent with the effect of K868R mutation that redirects the fate of AMPAR-containing vesicles from the degradative to the recycling pathway in neurons, thereby stabilising the number of AMPARs on the plasma membrane following neuronal activity (Schwarz et al., 2010; Widagdo et al., 2015). Our data also imply that a fraction of internalised GluA1-containing AMPARs is ubiquitinated and destined for lysosomal degradation, presumably to prevent excessive AMPARs from being inserted into the plasma membrane during LTP. Our findings also establish the requirement of GluA1 ubiquitination in mediating both NMDAR- and mGluR-dependent LTD. Although direct stimulation of primary neurons with NMDA and DHPG is not sufficient to induce GluA1 ubiquitination to a level detectable by western blotting, bicuculline-induced ubiquitination of GluA1 does involve NMDAR- and mGluR-dependent signalling (Widagdo et al., 2015).

GluA1 ubiquitination is not only important for LTP and LTD, but also contributes to spatial and short-term memory. In the novelty preference task, GluA1 K868R knock-in mice did not show any preference to explore the novel arm during the test session. This is in accordance with the essential role of GluA1 in short-term spatial memory (Bannerman et al., 2014; Sanderson et al., 2009). Using the active place avoidance behavioural task, we found that the GluA1 knock-in mice perform as well as their wild-type littermates, indicating that hippocampal-dependent spatial learning is intact in these ubiquitin-deficient mutant mice. These findings are consistent with the normal spatial learning reported in GluA1 knockout and S831/845A phospho-deficient mutant mice (Lee et al., 2003; Reisel et al., 2002; Zamanillo et al., 1999). However, we observed that reversal learning wa s impaired in the GluA1 ubiquitin-deficient mutant, which is associated with deficits in both NMDAR- and mGluR-mediated LTD. This is interesting considering that deficits in behavioural flexibility are often accompanied by defective GluA1 endocytosis and deficits in LTD (Awasthi et al., 2019; Collingridge et al., 2010; Eales et al., 2014; Mills et al., 2014; Nicholls et al., 2008). Taken together, our results indicate that the correct targeting and endosomal recycling of AMPARs to the postsynaptic membrane through the ubiquitination of GluA1 are necessary for the expression of rapidly learnt spatial memory, memory update and cognitive flexibility.

Several lines of evidence have demonstrated the direct involvement of AMPAR ubiquitination as a critical pathway in mediating Aβ-induced synaptic depression in neurons. Firstly, the expression of Nedd4-1 is upregulated in the human Alzheimer’s brain, which correlates with an increase in AMPAR ubiquitination and a decrease in total AMPAR protein levels (Kwak et al., 2012; Zhang et al., 2018). Secondly, knocking down the expression of Nedd4-1 abolishes the Aβ-induced reduction in the levels of surface AMPARs, AMPAR-mediated currents and dendritic spine loss (Rodrigues et al., 2016; Zhang et al., 2018). Thirdly, acute exposure of cultured neurons to soluble Aβ oligomers induces AMPAR ubiquitination concomitant with the removal of these receptors from the plasma membrane (Guntupalli et al., 2017). Importantly, the expression of GluA1-K868R ubiquitin-deficient mutants inhibits the adverse effects of Aβ on the surface expression of AMPARs in neurons. Therefore, the GluA1 K868R knock-in mice we have generated will be useful for determining whether inhibiting the ubiquitination of AMPARs is a viable strategy to prevent the effects of Aβ on cognitive functions in mouse models of Alzheimer’s disease.

## ACKNOWLEDGEMENTS

This work was supported by grants from the Australian Medical Research Future Fund (Clem Jones Centre for Ageing Dementia Research Flagship Project Grant) and the John T. Reid Charitable Trusts (to VA), and the National Honor Scientist Program (NRF-2012R1A3A1050385) of Korea (to BK). SG was a recipient of a UQ International Scholarship. JW was supported by a UQ Amplify award and an Australian Research Council DECRA Fellowship (DE170100112). We thank Dr Richard Huganir (Johns Hopkins University, Baltimore) for providing antibodies. We also thank Rowan Tweedale for editing this manuscript. Imaging was performed at the Queensland Brain Institute’s Advanced Microscopy Facility.

## AUTHOR CONTRIBUTIONS

VA conceived the project. DGB, DJJ, JW, BK and VA designed the research. SG, PP, DHH and MR performed experiments and analysed the data. FK contributed new reagents. BK and VA supervised the study. VA wrote the manuscript. All authors discussed the results and commented on the manuscript.

## STAR METHODS

### KEY RESOURCES TABLES

| REAGENT or RESOURCE | SOURCE | IDENTIFIER |
| --- | --- | --- |
| **Antibodies** |  |  |
| β-actin (rabbit monoclonal, clone 13E5, used 1:10,000 in western blotting) | Cell Signaling Technology | Cat# 4970, RRID:AB_2223172 |
| GluA1 (mouse monoclonal, clone 4.9D, used 1:1,000 in western blotting) | A gift from R. Huganir (Widagdo et al., 2015) | N/A |
| GluA2 (mouse monoclonal, clone 6A, used 1:1,000 in western blotting) | A gift from R. Huganir (Widagdo et al., 2015) | N/A |
| Peroxidase-conjugated donkey anti-rabbit IgG antibody | GE Healthcare | Cat# NA934, RRID:AB_772206 |
| Peroxidase-conjugated sheep anti-mouse IgG antibody | GE Healthcare | Cat# NA931, RRID:AB_772210 |
| PSD-95 (mouse polyclonal, clone K28/43, used 1:2,000 in western blotting) | NeuroMab | Cat# K28/43, RRID:AB_2877189 |
| Synaptophysin (mouse monoclonal, used 1:10,000 in western blotting) | Synaptic Systems | Cat# 101 011, RRID:AB_887824 |
| Ubiquitin (mouse polyclonal, clone P4D1, used 1:1,000 in western blotting) | Santa Cruz Biotechnology | Cat# sc-8017, RRID:AB_628423 |
| **Chemicals, Peptides, and Recombinant Proteins** |  |  |
| (S)-AMPA | Tocris | Cat# 0254/1 |
| D-AP5 sodium salt | Hello Bio | Cat# HB0225 |
| (+)-Bicuculline | Hello Bio | Cat# HB0896 |
| cOmplete, EDTA-free protease inhibitor cocktail | Sigma-Aldrich | Cat# 11873580001 |
| (R,S)-3,5-DHPG | Hello Bio | Cat# HB0026 |
| 5'-Fluoro-2'-deoxyuridine | Sigma-Aldrich | Cat# F0503 |
| N-ethylmaleimide | Sigma-Aldrich | Cat# E3876 |
| Papain suspension | Worthington | Cat# LS003126 |
| Paraformaldehyde | Sigma-Aldrich | Cat# P6148 |
| Picrotoxin | Hello Bio | Cat# HB0506 |
| Protein G-sepharose | GE Healthcare | Cat# GE17-0618-01 |
| Tetrodotoxin citrate | Abcam | Cat# ab120055 |
| Uridine | Sigma-Aldrich | Cat# U3003 |
| L-689,560 | Tocris | Cat# 0742 |
| **Experimental Models: Organisms/Strains** |  |  |
| Mouse: GluA1 K868R knock-in | This paper | N/A |
| **Software and Algorithms** |  |  |
| EthoVision XT | Noldus | RRID:SCR_000441; https://www.noldus.com/ethovision |
| Image Studio Lite | LI-COR | RRID:SCR_015795; https://www.licor.com/bio/image-studio-lite/ |
| GraphPad Prism 9 | GraphPad | RRID:SCR_002798; https://www.graphpad.com/ |
| Mini Analysis | Synaptosoft | RRID:SCR_002184; http://www.synaptosoft.com/MiniAnalysis |
| WinLTP | (Anderson and Collingridge, 2007) | https://www.winltp.com/index.html |

## RESOURCE AVAILABILITY

### Lead Contact

Further information and requests for resources and reagents should be directed to and will be fulfilled by the Lead contact, Victor Anggono.

### Material Availability

All unique reagents generated in this study are available from the Lead Contact with a completed Materials Transfer Agreement.

### Data and Code Availability

This study did not generate datasets/code.

## EXPERIMENTAL MODEL AND SUBJECT DETAILS

### Animals

All experimental procedures conducted on animals in this study were approved by the University of Queensland Animal Ethics Committee (QBI/047/18) and the Seoul National University Institutional Animal Care and Use Committee (SNU-130526-1-7). The GluA1 K868R knock-in mouse was generated using the conventional homologous recombination method. The targeting vector comprises an artificial wild-type exon 16, flanked by inverted LoxP2272 and FRT sites, followed by a neomycin cassette, an FRT site, an inversed exon 16 containing the Lys-868 substitution to arginine (K868R) and an inverted LoxP2272 flanked by a pair of inverted LoxP sites (Figure 1A). This construct was electroporated into C57Bl/6 embryonic stem cells. Positive clones were identified by polymerase chain reaction (PCR) and Southern blotting using the 5’, 3’ and neomycin probes. Positive clones were injected into C57Bl/6 blastocysts, and chimeras were crossed to OzFlp mice to remove the neo cassette from the germline. After confirmation of *neo* excision by Southern blot analysis, heterozygous conditional knock-in mice were backcrossed to C57Bl/6 animals to remove the *Flp* gene and the resulting conditional knock-in mice were bred to homozygosity. The *Flp*-negative mice were subsequently bred to CMV-Cre mice to obtain the GluA1 K868R knock-in heterozygous offspring. The *Cre* gene was then bred out by crossing these offspring to C57Bl/6 mice. The *Flp*- and *Cre*-negative GluA1 K868R knock-in mice were maintained as heterozygous mating.

### Primary cortical neuronal cultures

Cortices derived from embryonic day 17 mouse pups, either from wild-type or GluA1 K868R KI mice, were used to generate primary neuronal cultures (Widagdo et al., 2015). Cortices were isolated and dissociated with 30U of papain suspension (Worthington) for 20 min in a 37°C water bath. A single-cell suspension was obtained by triturating tissues with fire-polished glass Pasteur pipettes and then plated at a density of 1.2 × 10^6^ cells on poly-L-lysine-coated 6 cm tissue culture dishes in a Neurobasal growth medium (Invitrogen) supplemented with 2 mM Glutamax, 1% penicillin/streptomycin, 2% B27 and 5% foetal calf serum (FBS). Cortical neurons were maintained in a Neurobasal medium containing 1% FBS and fed twice a week. Uridine (Sigma, 5 μM) and 5’-fluoro-2’-deoxyuridine (Sigma, 5 μM) were added to the culture medium at days *in vitro* (DIV) 5 to stop glial proliferation. Cells were processed at DIV 14-16.

## METHOD DETAILS

### Genotyping

Genotyping of GluA1 KI mice was carried out by PCR of genomic DNA isolated from ear or toe biopsies using three primers: primer 1 (5’-ATGGCAGAGTGGTCAGCCAG-3’), primer 2 (5’-TTTGGCACTGAAGGGCTTGG-3’) and primer 3 (5’-GCTATCCGGACCAGTACACT-3’). The wild-type allele produces a band of 172 base pairs (bp), whereas the knock-in allele yields a 222-bp product when resolved on a 3% agarose gel. PCR fragments were sent for sequencing to validate the presence of KI mutation.

### Subcellular fractionation

Forebrains were isolated from mice and homogenised in ice-cold sucrose buffer A (0.32 M sucrose, 10 mM HEPES, pH 7.4). The homogenate was centrifuged at 1,000 *g* for 10 min at 4°C, yielding the supernatant fraction and the nuclear-enriched pellet. An aliquot of the homogenate was taken for further analysis as the total protein fraction, with the remainder being centrifuged at 10,000 *g* for 15 min at 4°C to obtain a crude synaptosomal (P2) fraction and cytosolic protein-enriched supernatant (S2). The P2 fraction was resuspended with sucrose buffer and centrifuged at 10,000 *g* for 15 min at 4°C. The resulting pellets were lysed by hypo-osmotic shock in ice-cold water and homogenised by pipetting. One M HEPES was rapidly added to a final concentration of 4 mM to adjust the pH. The P2 suspension was incubated by rotating in the cold room for 30 min before an ultracentrifugation step at 25,000 *g* for 20 min. The resulting pellet (LP1) was further resuspended in buffer B (50 mM HEPES, 2 mM EDTA, pH. 7.4) containing 1% Triton X-100 and mixed in a cold room for 15 min. This LP1 suspension was then centrifuged at 32,000 *g* for 20 min to obtain the PSD pellet. All fractions were denatured after adjusting total protein concentrations followed by western blotting.

### Ubiquitination assay

Ubiquitination of GluA1 and GluA2 was induced by incubating neurons in artificial cerebrospinal fluid (ACSF) containing 100 μM AMPA plus 50 μM D-APV and 1 μM tetrodotoxin (TTX) for 10 min at 37°C (Widagdo et al., 2015). Neurons were lysed in warm 1% SDS (in phosphate-buffered saline [PBS]) and diluted in ten volumes of ice-cold cell lysis buffer (1% Triton X-100, 1 mM EDTA, 1 mM EGTA, 50 mM NaF, and 5 mM Na-pyro-phosphate in PBS) supplemented with 10 mM *N*-ethylmaleimide and Complete EDTA-free protease inhibitor cocktails (Sigma). Lysates were centrifuged at 14,000 rpm for 20 min at 4°C and cleared with protein G-sepharose beads. Pre-cleared lysates were then incubated with GluA1 or GluA2 antibodies coupled to protein G-Sepharose overnight at 4°C, followed by four washes with ice-cold lysis buffer and elution in 2X SDS sample buffer. The immunoprecipitated proteins were resolved by SDS-PAGE and probed by western blot analysis with specific antibodies against GluA1, GluA2 and ubiquitin.

### Western blot

Protein samples were resolved on 6%, 10% or 15% SDS polyacrylamide gels and transferred to a PVDF (polyvinylidene fluoride) membrane (Millipore) at 100 V for 2 h. Membranes were blocked with 5% skim milk in Tris-buffered saline (TBS) containing 0.01% Tween-20 for 1 h and incubated with primary antibody overnight at 4°C. After extensive washing with 1% skim milk, horseradish peroxidase (HRP)-conjugated secondary antibodies were added and incubated for 1 h at room temperature under constant shaking. Signals were developed with an enhanced chemiluminescence (ECL) method. Images were acquired on the Odyssey Fc imaging system (LI-COR) and band intensities were quantified using Image Studio Lite software (LI-COR).

### Nissl staining

Mice were deeply anaesthetised and perfused transcardially with phosphate-buffered saline (PBS), followed by a 4% paraformaldehyde (PFA) solution. Dissected brains were immersed in 4% PFA for 24 h. They were then washed three times with distilled water, immersed with PBS containing sodium azide as a preservative, and stored at 4°C until used. Brains were sliced coronally using a vibratome at a thickness of 50 µm. Sections were mounted on glass slides and stained with 0.1% cresyl violet for 5 min. Images were taken using a scanning bright-field microscope.

### Electrophysiological recordings

Adult male mice were used for all electrophysiological recordings (2-5 months of age). Animals were deeply anaesthetised with isoflurane and sacrificed by decapitation. Transverse hippocampal slices (350 μm) were prepared using a vibratome (Leica, VT1200S) in an ice-chilled slicing solution that contained (in mM): 210 sucrose, 3 KCl, 26 NaHCO_3_, 1.25 NaH_2_PO_4_, 5 MgSO_4_, 10 D-glucose, 3 sodium ascorbate and 0.5 CaCl_2_, saturated with 95% O_2_ and 5% CO_2_. The slices were transferred to an incubation chamber that contained the recording solution (ACSF; mM): 124 NaCl, 3 KCl, 26 NaHCO_3_, 1.25 NaH_2_PO_4_, 2 MgSO_4_, 10 D-glucose and 2 CaCl_2_ (carbonated with 95% O_2_ and 5% CO_2_). Slices were allowed to recover at 32-34°C for 30 min and then maintained at 26-28 °C for a minimum of 1 h before recordings were made.

Standard extracellular recordings were performed in the CA1 region of the hippocampal slices maintained at 32°C, as described previously (Park et al., 2016), to measure the slope of the evoked field excitatory postsynaptic potentials (fEPSPs). Responses were obtained using Multiclamp 700B amplifier (Molecular Devices) and digitised with a Digidata 1322A A/D board at a sampling rate of 20 kHz (low-pass filtered at 10 kHz). Recordings were monitored and analysed using WinLTP (Anderson and Collingridge, 2007). Schaffer collateral-commissural pathways were stimulated at a baseline frequency of 0.1 Hz. After a stable baseline of at least 20 min, LTP was induced using theta-burst stimulation (TBS) delivered at the basal stimulus intensity. An episode of TBS comprised five bursts at 5 Hz, with each burst composed of five pulses at 100 Hz. Either a single episode of TBS or a train of three TBS episodes at an inter-episode interval of 10 min was given. LTD was induced using a repetitive low-frequency stimulation (LFS) at 1 Hz for 15 min, where P14 mice were used for a reliable LFS-LTD induction. DHPG-LTD was induced by the bath application of DHPG (100 µM) in the presence of D-AP5 (50 µM) to prevent non-specific NMDAR-mediated effects. Representative sample traces were an average of four consecutive responses, collected from typical experiments (stimulus artefacts were blanked for clarity). The input-output relationships were estimated by varying the stimulus intensity from 1.5 to 15 V in 1.5 V increments. Paired-pulse facilitation was ranged at inter-stimulus intervals from 20 to 400 ms at the basal stimulus intensity.

Whole-cell recording was performed at 32°C during continuous perfusion at 3-4 ml/min with ACSF that contained 50 μM picrotoxin and 10 µM (+)-bicuculline to prevent GABA_A_ receptor-mediated transmission. CA1 pyramidal cells were visualised with IR-DIC optics (Olympus). The whole-cell solution comprised (mM): 8 NaCl, 130 CsMeSO_3_, 10 HEPES, 0.5 EGTA, 4 Mg-ATP, 0.3 Na_3_-GTP, 5 QX-314 and 0.1 spermine. The pH was adjusted to 7.2-7.3 with CsOH and osmolarity was set to 285-290 mOsm/l. The Schaffer collateral-commissural pathway was stimulated at a constant frequency of 0.1 Hz. Borosilicate glass pipettes were used with a resistance of 4-6 MΩ, and experiments were only accepted for analysis if series resistance values were <25 MΩ and varied by <20% during the course of the experiment. Signals were filtered at 10 kHz and digitised at 20 kHz using a Multiclamp 700B (Molecular Devices). Evoked excitatory postsynaptic currents (EPSCs) were monitored and analysed using WinLTP. For NMDA/AMPA ratio, cells were clamped at a holding potential of -70 mV to measure the peak of AMPAR-mediated synaptic transmission. NMDAR-currents were estimated at 50 ms after the stimulation onset at +40 mV of holding potential. Averages of 15 consecutive responses obtained at these holding potentials were used for the ratio. For rectification index (RI) measurements, AMPAR-mediated currents were pharmacologically isolated using a combination of a competitive NMDAR antagonist (D-AP5; 50 µM) plus a glycine-site antagonist (L-689,560; 5 µM). Neurons were immediately depolarised to +40 mV for 100 s, then to 0 mV for 50 s, and consecutive responses obtained at these holding potentials were averaged and calculated by taking the ratio of the slopes from 0 to +40 mV and -70 to 0 mV. The amplitude and frequency of miniature EPSCs were analysed using Mini Analysis (Synaptosoft) from the data obtained for 3 min at a holding potential of -70 mV in the presence of TTX (0.5 µM).

### Animal handling

Mice were maintained under standard conditions on a 12-h light/dark cycle (lights on at 7:00 am) with food and water provided *ad libitum*. All experiments were conducted between 9:00 am and 5 pm. The mice were group-housed with 2-5 animals per cage. Approximately 12- to 15-week-old littermate pairs of wild-type and KI male mice were used for all experiments. Mice were handled for 3 min per day for 3 consecutive days before the commencement of experiments.

### Active place avoidance test

The active place avoidance task is a hippocampal-dependent spatial learning test for rodents (Willis et al., 2017). This task was performed on an elevated rotating circular arena (at 1 rotation per min) located in a room with visual cues on the walls. The arena contained a 60º shock zone with an electric grid that delivered a brief foot shock (0.5 mA for 500 ms) upon animal entry. A second foot shock was delivered at 1.5 s intervals if the animal remained in the shock zone. The position of the shock zone was kept constant relative to the spatial frame of the room. One day after habituation, mice were tested for 10 min in each training session over 5 days. The position of the animal was tracked using an overhead camera linked to Tracker software (Bio-Signal group). To assess the cognitive flexibility of mice, we also performed a reversal test on day 6, in which the position of the shock zone was moved opposite to the initial location. Mice were tested for 10 min in each session for a further 5 days. The number of entries into the shock zone and the distance travelled were measured to determine the extent of learning.

### Y maze

A Y maze with a blocked arm was used to measure short-term spatial memory in mice. The apparatus comprised three arms made of Perspex elevated above the floor and was placed under an overhead camera. Lighting conditions (400 lux) were kept constant throughout the experiment. In the first trial, mice were placed in the starting arm and allowed to explore the open arm for 5 min. The second trial began 90 min later in which mice were placed back into the starting arm of the maze with the Perspex divider removed. Mice were allowed to explore all arms for 5 min. Recorded videos were analysed using Ethovision XT9 image analysis software (Noldus). The number of entries and time spent in the starting and novel arms were determined for each animal.

### Elevated plus maze

An elevated plus maze was used to determine anxiety-related behaviour. The apparatus was a plus-shaped opaque Perspex elevated platform with two open (64.5 × 6.2 cm) and two enclosed arms (64.5 × 6.2 cm) radiating from the centre platform. The apparatus was placed under an overhead camera to track the animal’s movement. Lighting conditions (400 lux) were kept constant throughout the experiment. Mice were placed in the middle of the maze and allowed to explore for 5 min. Recorded videos were analysed using Ethovision XT9 image analysis software. The number of entries and time spent in the closed and open arms were determined for each animal.

### Open field test

The open field test was used to assay locomotor activity and anxiety-like behaviour. This test was conducted in a square arena (30.7 × 30.7 × 30.7 cm) placed underneath an overhead camera which tracked the animal’s movement. Lighting conditions (400 lux) were kept constant throughout the experiment. In an individual experiment, each mouse was placed in the centre of the arena and allowed to explore for 3 min. Recorded videos were analysed using Ethovision XT9 image analysis software. The total distance travelled was measured for the duration of the trial, together with the time spent in and the number of entries into the centre of the arena.

### Statistical analysis

All statistical analyses were performed using Prism8 software (GraphPad Software). Comparison between individual experimental groups was performed by unpaired Student’s t-test where appropriate. Comparison of more than two parameters/multiple groups was performed by two-way ANOVA with Sidak’s multiple comparison test. All graphs were presented as mean ± SEM. The difference between means from experimental groups was considered significant at the level of *p* < 0.05.

## SUPPLEMENTARY FIGURES

**Supplementary Figure 1.**
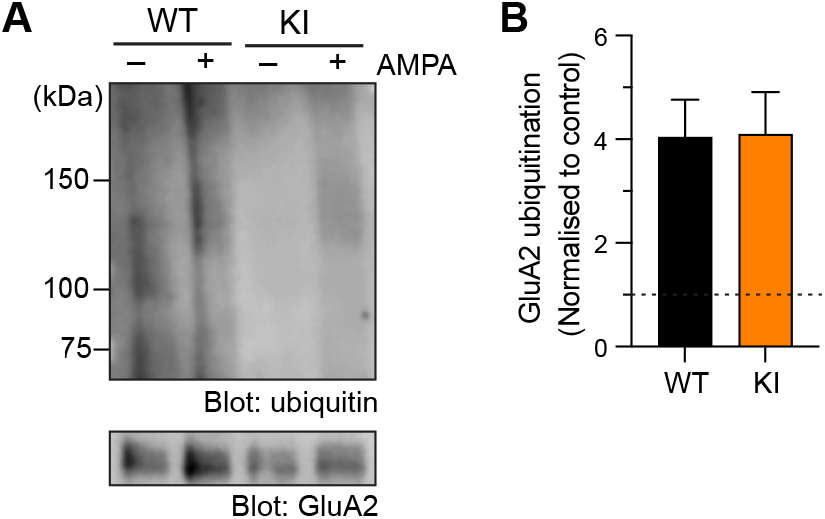
related to Figure 1. Normal AMPA-induced GluA2 ubiquitination in neurons derived from GluA1 K868R knock-in mice. (A) Primary cortical neurons derived from wild-type (WT) or GluA1 K868R knock-in (KI) mice were incubated in the presence or absence of 100 μM AMPA, lysed and immunoprecipitated with anti-GluA1 antibodies. Eluted proteins were subjected to western blot analysis and probed with anti-ubiquitin and anti-GluA1 antibodies. (B) Quantification of the levels of AMPA-induced GluA2 ubiquitination normalised to the controls (dashed line). Data are presented as mean ± SEM (n = 6 per group). *p* > 0.99 using a Mann-Whitney test.

**Supplementary Figure 2.**
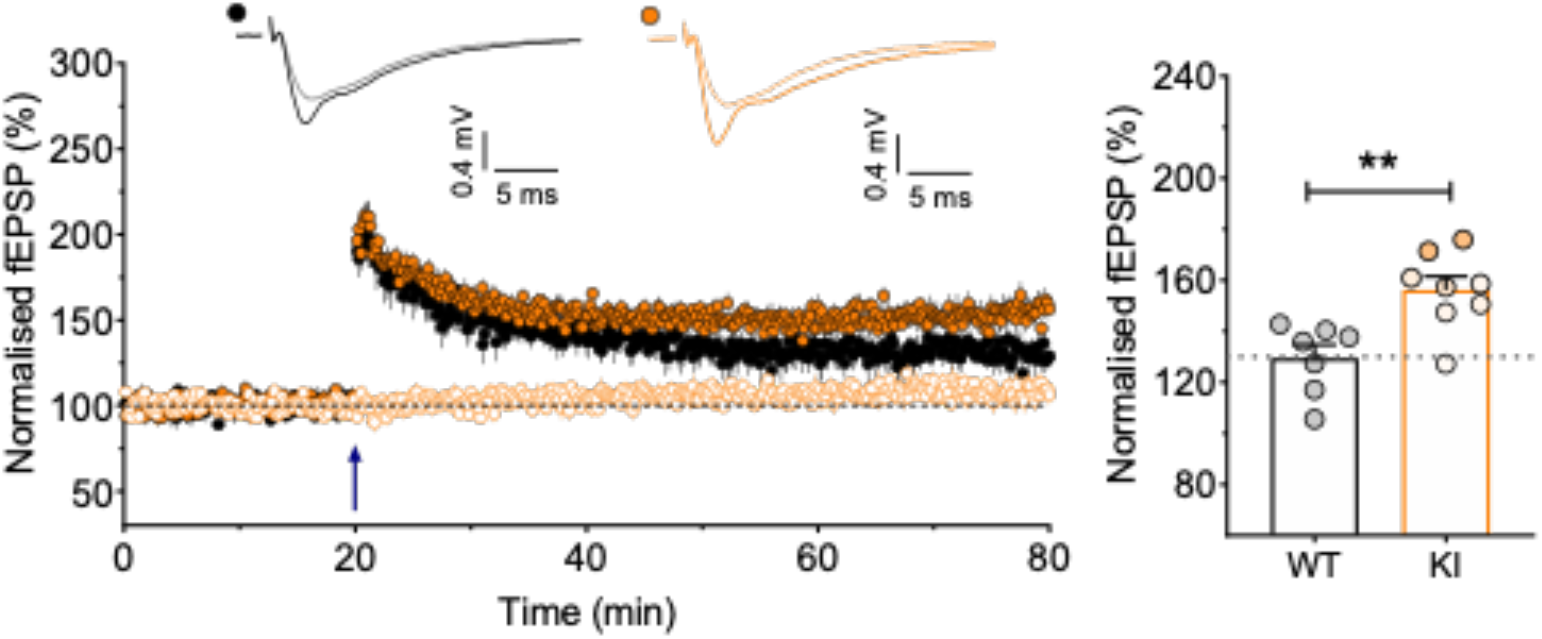
related to Figure 2. Enhanced LTP in GluA1 K868R mutant mice. Input-specific LTP induced by a single episode of theta-burst stimulation (5 bursts at 5 Hz, with each burst composed of five pulses at 100 Hz). Quantification for the levels of LTP in WT (n = 7 from 7 mice) and KI hippocampal slices (n = 8/8). \*\**p* = 0.004 using an unpaired t-test.

**Supplementary Figure 3.**
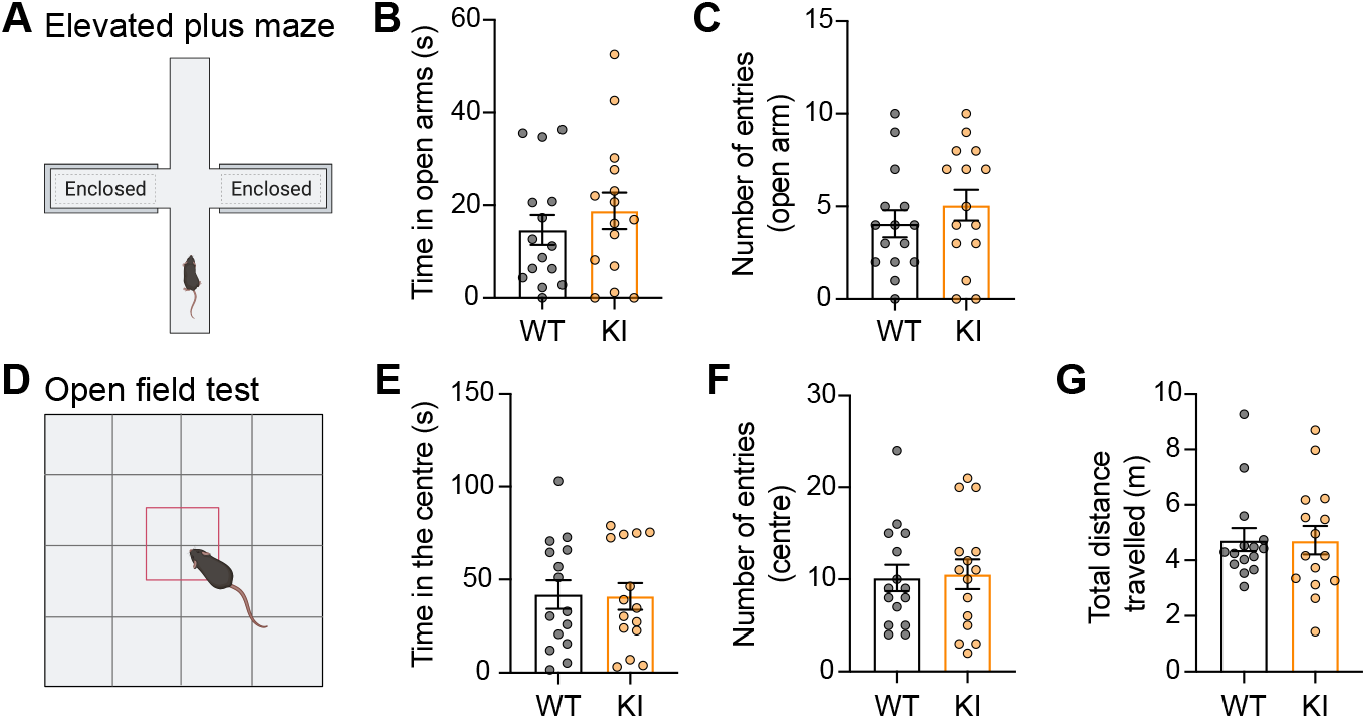
related to Figure 4. GluA1 K868R knock-in mice display normal locomotor activity and anxiety-like behaviour. (A – C) Measurement of anxiety-like behaviour using the elevated plus maze (A) revealed no differences in the time spent (B) or the number of entries (C) to the open arms between genotypes. Data are presented as mean ± SEM (WT, n = 15 and KI, n = 15). *p =* 0.486 (B) and *p* = 0.380 (C) using Mann-Whitney tests. (D – G) Measurement of locomotor activity and anxiety-like behaviour using the open field test (D) revealed no differences in the time spent (E), the number of entries (F) in to the centre zone, or the total distance travelled (G) between genotypes. Data are presented as mean ± SEM (WT, n = 15 and KI, n = 15). *p =* 0.806 (E), *p* = 0.91 (F) and p = 0.87 (G) using Mann-Whitney tests.

**Supplementary Data 1.**
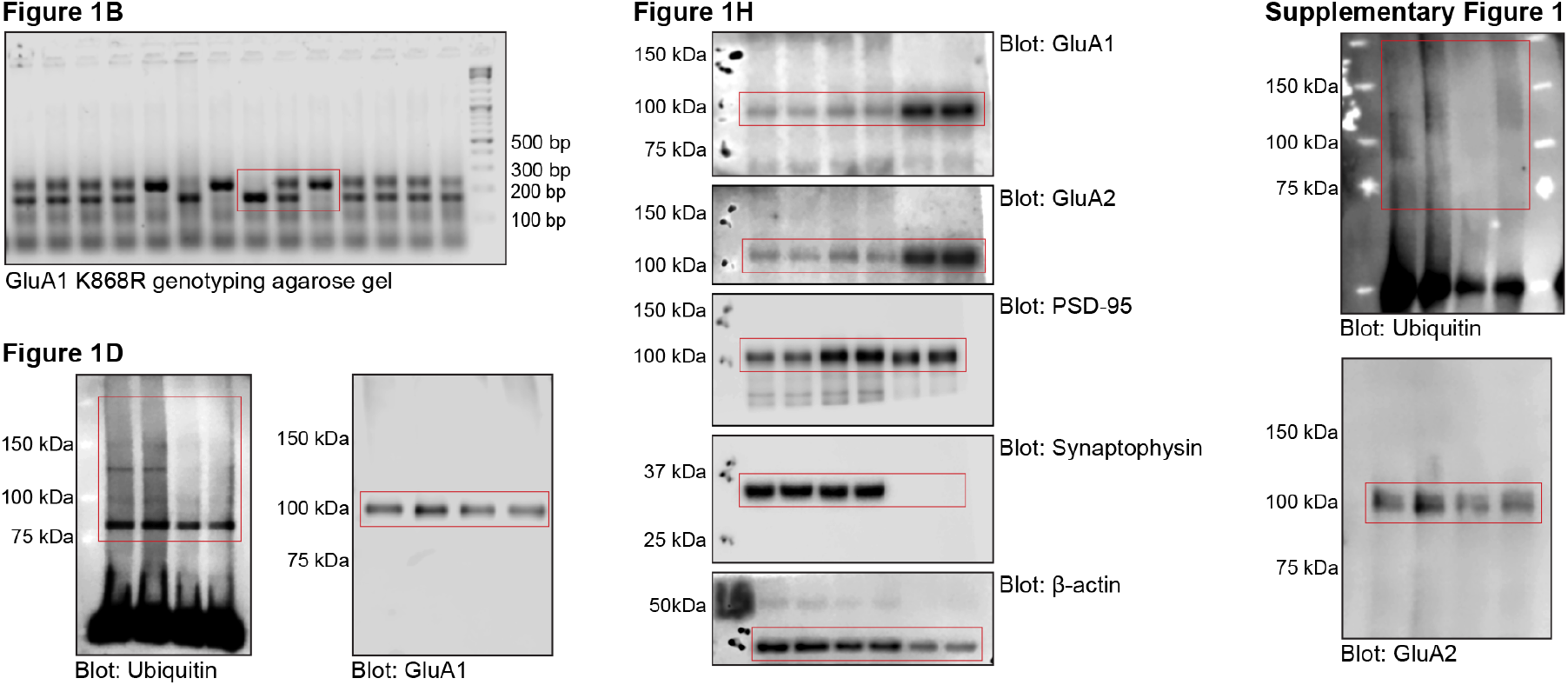
Raw images for agarose gel and western blots.

## REFERENCES

Anderson, W.W., and Collingridge, G.L. (2007). Capabilities of the WinLTP data acquisition program extending beyond basic LTP experimental functions. J Neurosci Methods 162, 346–356.

Anggono, V., and Huganir, R.L. (2012). Regulation of AMPA receptor trafficking and synaptic plasticity. Curr Opin Neurobiol 22, 461–469.

Awasthi, A., Ramachandran, B., Ahmed, S., Benito, E., Shinoda, Y., Nitzan, N., Heukamp, A., Rannio, S., Martens, H., Barth, J., et al. (2019). Synaptotagmin-3 drives AMPA receptor endocytosis, depression of synapse strength, and forgetting. Science 363, eaav1483.

Bannerman, D.M., Sprengel, R., Sanderson, D.J., McHugh, S.B., Rawlins, J.N., Monyer, H., and Seeburg, P.H. (2014). Hippocampal synaptic plasticity, spatial memory and anxiety. Nat Rev Neurosci 15, 181–192.

Brown, T.C., Correia, S.S., Petrok, C.N., and Esteban, J.A. (2007). Functional compartmentalization of endosomal trafficking for the synaptic delivery of AMPA receptors during long-term potentiation. J Neurosci 27, 13311–13315.

Collingridge, G.L., Peineau, S., Howland, J.G., and Wang, Y.T. (2010). Long-term depression in the CNS. Nat Rev Neurosci 11, 459–473.

Diering, G.H., and Huganir, R.L. (2018). The AMPA receptor code of synaptic plasticity. Neuron 100, 314–329.

Eales, K.L., Palygin, O., O’Loughlin, T., Rasooli-Nejad, S., Gaestel, M., Muller, J., Collins, D.R., Pankratov, Y., and Correa, S.A. (2014). The MK2/3 cascade regulates AMPAR trafficking and cognitive flexibility. Nat Commun 5, 4701.

Fernandez-Monreal, M., Brown, T.C., Royo, M., and Esteban, J.A. (2012). The balance between receptor recycling and trafficking toward lysosomes determines synaptic strength during long-term depression. J Neurosci 32, 13200–13205.

Goo, M.S., Scudder, S.L., and Patrick, G.N. (2015). Ubiquitin-dependent trafficking and turnover of ionotropic glutamate receptors. Front Mol Neurosci 8, 60.

Guntupalli, S., Jang, S.E., Zhu, T., Huganir, R.L., Widagdo, J., and Anggono, V. (2017). GluA1 subunit ubiquitination mediates amyloid-β-induced loss of surface α-amino-3-hydroxy-5-methyl-4-isoxazolepropionic acid (AMPA) receptors. J Biol Chem 292, 8186–8194.

Hayashi, T. (2021). Post-translational palmitoylation of ionotropic glutamate receptors in excitatory synaptic functions. Br J Pharmacol 178, 784–797.

Hu, H., Real, E., Takamiya, K., Kang, M.G., Ledoux, J., Huganir, R.L., and Malinow, R. (2007). Emotion enhances learning via norepinephrine regulation of AMPA-receptor trafficking. Cell 131, 160–173.

Huganir, R.L., and Nicoll, R.A. (2013). AMPARs and synaptic plasticity: the last 25 years. Neuron 80, 704–717.

Kraeuter, A.K., Guest, P.C., and Sarnyai, Z. (2019). The Y-maze for assessment of spatial working and reference memory in mice. Methods Mol Biol 1916, 105–111.

Kristensen, A.S., Jenkins, M.A., Banke, T.G., Schousboe, A., Makino, Y., Johnson, R.C., Huganir, R., and Traynelis, S.F. (2011). Mechanism of Ca2+/calmodulin-dependent kinase II regulation of AMPA receptor gating. Nat Neurosci 14, 727–735.

Kwak, Y.D., Wang, B., Li, J.J., Wang, R., Deng, Q., Diao, S., Chen, Y., Xu, R., Masliah, E., Xu, H., et al. (2012). Upregulation of the E3 ligase NEDD4-1 by oxidative stress degrades IGF-1 receptor protein in neurodegeneration. J Neurosci 32, 10971–10981.

Lee, H.K., Barbarosie, M., Kameyama, K., Bear, M.F., and Huganir, R.L. (2000). Regulation of distinct AMPA receptor phosphorylation sites during bidirectional synaptic plasticity. Nature 405, 955–959.

Lee, H.K., Takamiya, K., Han, J.S., Man, H., Kim, C.H., Rumbaugh, G., Yu, S., Ding, L., He, C., Petralia, R.S., et al. (2003). Phosphorylation of the AMPA receptor GluR1 subunit is required for synaptic plasticity and retention of spatial memory. Cell 112, 631–643.

Lin, A., Hou, Q., Jarzylo, L., Amato, S., Gilbert, J., Shang, F., and Man, H.Y. (2011). Nedd4-mediated AMPA receptor ubiquitination regulates receptor turnover and trafficking. J Neurochem 119, 27–39.

Lu, W., and Roche, K.W. (2012). Posttranslational regulation of AMPA receptor trafficking and function. Curr Opin Neurobiol 22, 470–479.

Lu, W., Shi, Y., Jackson, A.C., Bjorgan, K., During, M.J., Sprengel, R., Seeburg, P.H., and Nicoll, R.A. (2009). Subunit composition of synaptic AMPA receptors revealed by a single-cell genetic approach. Neuron 62, 254–268.

Lussier, M.P., Sanz-Clemente, A., and Roche, K.W. (2015). Dynamic regulation of N-methyl-d-aspartate (NMDA) and α-amino-3-hydroxy-5-methyl-4-isoxazolepropionic acid (AMPA) receptors by posttranslational modifications. J Biol Chem 290, 28596–28603.

Mabb, A.M. (2021). Historical perspective and progress on protein ubiquitination at glutamatergic synapses. Neuropharmacology 196, 108690.

Makino, Y., Johnson, R.C., Yu, Y., Takamiya, K., and Huganir, R.L. (2011). Enhanced synaptic plasticity in mice with phosphomimetic mutation of the GluA1 AMPA receptor. Proc Natl Acad Sci U S A 108, 8450–8455.

Man, H.Y., Sekine-Aizawa, Y., and Huganir, R.L. (2007). Regulation of α-amino-3-hydroxy-5-methyl-4-isoxazolepropionic acid receptor trafficking through PKA phosphorylation of the Glu receptor 1 subunit. Proc Natl Acad Sci U S A 104, 3579–3584.

Mills, F., Bartlett, T.E., Dissing-Olesen, L., Wisniewska, M.B., Kuznicki, J., Macvicar, B.A., Wang, Y.T., and Bamji, S.X. (2014). Cognitive flexibility and long-term depression (LTD) are impaired following betacatenin stabilization in vivo. Proc Natl Acad Sci U S A 111, 8631–8636.

Nicholls, R.E., Alarcon, J.M., Malleret, G., Carroll, R.C., Grody, M., Vronskaya, S., and Kandel, E.R. (2008). Transgenic mice lacking NMDAR-dependent LTD exhibit deficits in behavioral flexibility. Neuron 58, 104–117.

Park, M., Penick, E.C., Edwards, J.G., Kauer, J.A., and Ehlers, M.D. (2004). Recycling endosomes supply AMPA receptors for LTP. Science 305, 1972–1975.

Park, P., Sanderson, T.M., Amici, M., Choi, S.L., Bortolotto, Z.A., Zhuo, M., Kaang, B.K., and Collingridge, G.L. (2016). Calcium-permeable AMPA receptors mediate the induction of the protein kinase A-dependent component of long-term potentiation in the hippocampus. J Neurosci 36, 622–631.

Park, P., Volianskis, A., Sanderson, T.M., Bortolotto, Z.A., Jane, D.E., Zhuo, M., Kaang, B.K., and Collingridge, G.L. (2014). NMDA receptor-dependent long-term potentiation comprises a family of temporally overlapping forms of synaptic plasticity that are induced by different patterns of stimulation. Philos Trans R Soc Lond B Biol Sci 369, 20130131.

Parkinson, G.T., and Hanley, J.G. (2018). Mechanisms of AMPA receptor endosomal sorting. Front Mol Neurosci 11, 440.

Petrini, E.M., Lu, J., Cognet, L., Lounis, B., Ehlers, M.D., and Choquet, D. (2009). Endocytic trafficking and recycling maintain a pool of mobile surface AMPA receptors required for synaptic potentiation. Neuron 63, 92–105.

Reisel, D., Bannerman, D.M., Schmitt, W.B., Deacon, R.M., Flint, J., Borchardt, T., Seeburg, P.H., and Rawlins, J.N. (2002). Spatial memory dissociations in mice lacking GluR1. Nat Neurosci 5, 868–873.

Rodrigues, E.M., Scudder, S.L., Goo, M.S., and Patrick, G.N. (2016). Aβ-induced synaptic alterations require the E3 ubiquitin ligase Nedd4-1. J Neurosci 36, 1590–1595.

Sanderson, D.J., Good, M.A., Skelton, K., Sprengel, R., Seeburg, P.H., Rawlins, J.N., and Bannerman, D.M. (2009). Enhanced long-term and impaired short-term spatial memory in GluA1 AMPA receptor subunit knockout mice: evidence for a dual-process memory model. Learn Mem 16, 379–386.

Sanderson, D.J., Gray, A., Simon, A., Taylor, A.M., Deacon, R.M., Seeburg, P.H., Sprengel, R., Good, M.A., Rawlins, J.N., and Bannerman, D.M. (2007). Deletion of glutamate receptor-A (GluR-A) AMPA receptor subunits impairs one-trial spatial memory. Behav Neurosci 121, 559–569.

Sanderson, D.J., McHugh, S.B., Good, M.A., Sprengel, R., Seeburg, P.H., Rawlins, J.N., and Bannerman, D.M. (2010). Spatial working memory deficits in GluA1 AMPA receptor subunit knockout mice reflect impaired short-term habituation: evidence for Wagner’s dual-process memory model. Neuropsychologia 48, 2303–2315.

Schwarz, L.A., Hall, B.J., and Patrick, G.N. (2010). Activity-dependent ubiquitination of GluA1 mediates a distinct AMPA receptor endocytosis and sorting pathway. J Neurosci 30, 16718–16729.

Widagdo, J., Chai, Y.J., Ridder, M.C., Chau, Y.Q., Johnson, R.C., Sah, P., Huganir, R.L., and Anggono, V. (2015). Activity-dependent ubiquitination of GluA1 and GluA2 regulates AMPA receptor intracellular sorting and degradation. Cell Rep 10, 783–795.

Widagdo, J., Guntupalli, S., Jang, S.E., and Anggono, V. (2017). Regulation of AMPA receptor trafficking by protein ubiquitination. Front Mol Neurosci 10, 347.

Widagdo, J., Kerk, J.W., Guntupalli, S., Huganir, R.L., and Anggono, V. (2020). Subunit-specific augmentation of AMPA receptor ubiquitination by phorbol ester. Cell Mol Neurobiol 40, 1213–1222.

Willis, E.F., Bartlett, P.F., and Vukovic, J. (2017). Protocol for short- and longer-term spatial learning and memory in mice. Front Behav Neurosci 11, 197.

Zamanillo, D., Sprengel, R., Hvalby, O., Jensen, V., Burnashev, N., Rozov, A., Kaiser, K.M., Koster, H.J., Borchardt, T., Worley, P., et al. (1999). Importance of AMPA receptors for hippocampal synaptic plasticity but not for spatial learning. Science 284, 1805–1811.

Zhang, Y., Guo, O., Huo, Y., Wang, G., and Man, H.Y. (2018). Amyloid-β induces AMPA receptor ubiquitination and degradation in primary neurons and human brains of Alzheimer’s disease. J Alzheimers Dis 62, 1789–1801.

Zhu, J., Lee, K.Y., Jewett, K.A., Man, H.Y., Chung, H.J., and Tsai, N.P. (2017). Epilepsy-associated gene Nedd4-2 mediates neuronal activity and seizure susceptibility through AMPA receptors. PLoS Genet 13, e1006634.

